# Refractive index mapping below the diffraction limit via single molecule localization microscopy

**DOI:** 10.1101/2025.08.20.670782

**Authors:** Simon Jaritz, Lukas Velas, Anna Gaugutz, Manuel Rufin, Philipp J. Thurner, Orestis G. Andriotis, Julian G. Maloberti, Simon Moser, Alexander Jesacher, Gerhard J. Schütz

## Abstract

Single molecule localization microscopy (SMLM) is a powerful method to image biological samples in three dimensions below the diffraction limit of light microscopy. Beyond the position of the emitter, the shape of the single molecule point spread function provides additional information, for example about the refractive properties of the sample between the emitter and the glass coverslip. Here, we show that combination of SMLM with atomic force microscopy (AFM) allows to map the refractive index of a biological sample at sub-diffraction resolution and at a precision only limited by measurement errors of SMLM and AFM. We showcase the new method by the determination of the refractive index of isolated single collagen fibrils. Variabilities both in refractive index and the swelling behavior of single fibrils upon drying and rehydration exposed deviations from the ensemble behavior, demonstrating differential hydration of single collagen fibrils. Mapping the refractive index along single collagen fibrils revealed substantial fluctuations at characteristic length scales below 500 nm, which indicates structural heterogeneity of collagen fibrils at the length scale of single collagen molecules.

## Introduction

Single molecule localization microscopy has boosted our information on the spatial organization of biological samples, especially cells ^1^. The overall idea is to employ rare stochastic appearances of single molecule signals in order to separate them from those of neighboring molecules, which allows circumventing the diffraction limit of light microscopy. Appearances can be due to the stochastic blinking of dye molecules in STORM or PALM, or the stochastic binding of fluorescently labelled ligands in PAINT. The obtained signals then allow for determining the 2D center of mass as proxy for the 2D position with a precision that is mainly limited by the signal to noise ratio. At high signal quality, localization precisions below 1 nm have been reported ^2^.

Also, the shape of the single molecule point spread function (psf) contains valuable information. For example, its width allows for extracting a dye’s position along the optical axis ^3,4^. This can further be improved by introducing artificial astigmatism ^5,6^ or other phase-shaping methods ^7^.

We have recently introduced defocused imaging in combination with supercritical angle detection, which allows for achieving impressive localization precision down to 10 nm in case emitters are located close to surfaces of high refractive index ^8,9^. In addition, the psf shape also varies with dipole angle, which has been used to determine azimuth and inclination angle precisely ^10^.

The above-mentioned research problems are all well-conditioned in a sense that the input argument’s variation has a distinct effect on the psf shape. In contrast, there are also ill-conditioned problems, where the variations of different parameters have nearly identical effects on the psf shape. One example is the refractive index *n* of the sample, which affects the psf similarly as the distance from the focal plane. Therefore, if *n* was not considered correctly the calculated axial distances will be distorted compared to the true distances ^11^. Conversely, psf analysis could also be used to determine the refractive index of the sample if the axial distance was known.

We reasoned that a bimodal imaging approach, in which SMLM is combined with a second imaging modality to record the axial position of the dyes, would transform the ill-conditioned into a well-conditioned problem for refractive index mapping at a resolution below the diffraction limit of light microscopy. For proof-of-principle, we used here a combination of SMLM with atomic force microscopy (AFM) to determine the refractive index of collagen fibrils mounted onto glass coverslips. For the achieved signal to noise ratio, the method allows for determining the refractive index to a precision of *σ*_*n*_ < 3 · 10^−3^. We applied the method to compare the refractive index of differently swollen collagen fibrils: with increasing water content we observed the refractive index of single collagen fibrils to decrease towards the refractive index of water.

## Results

The concept of super-resolved refractive index mapping is sketched in **Fig. 1a**. A fluorescently labelled sample with refractive index *n*_*s*_ is mounted on a glass coverslip with refractive index *n*_*g*_ and imaged both with AFM and SMLM. AFM provides the ground truth information on the distance of the fluorophores from the coverslip surface, *H*_*AFM*_. Next, SMLM is performed on the very same sample. **Fig. 1b** shows two hypothetical psfs calculated for a distance of *H* = 100 nm from the coverslip surface, which differ only in the refractive index of the underlying material: here, we compared *n*_*s*_ = 1.35 with *n*_*s*_ = 1.48. The difference image in **Fig. 1c** shows a broadened psf for the higher refractive index. Without independent information on the sample’s refractive index, one would hence wrongly estimate the fluorophore’s distance from the coverslip surface, *H*_*SMLM*_(*n*_*s*_): erroneous assumption of a refractive index smaller than the correct value would yield an underestimation of the molecule’s distance, and vice versa. *n*_*s*_ can therefore be derived by variation in the molecule image model to achieve an optimal fit to the observations. This is possible because the molecules’ distance to the coverslip is known from independent AFM measurements, *H*_*AFM*_.

**Figure 1.**
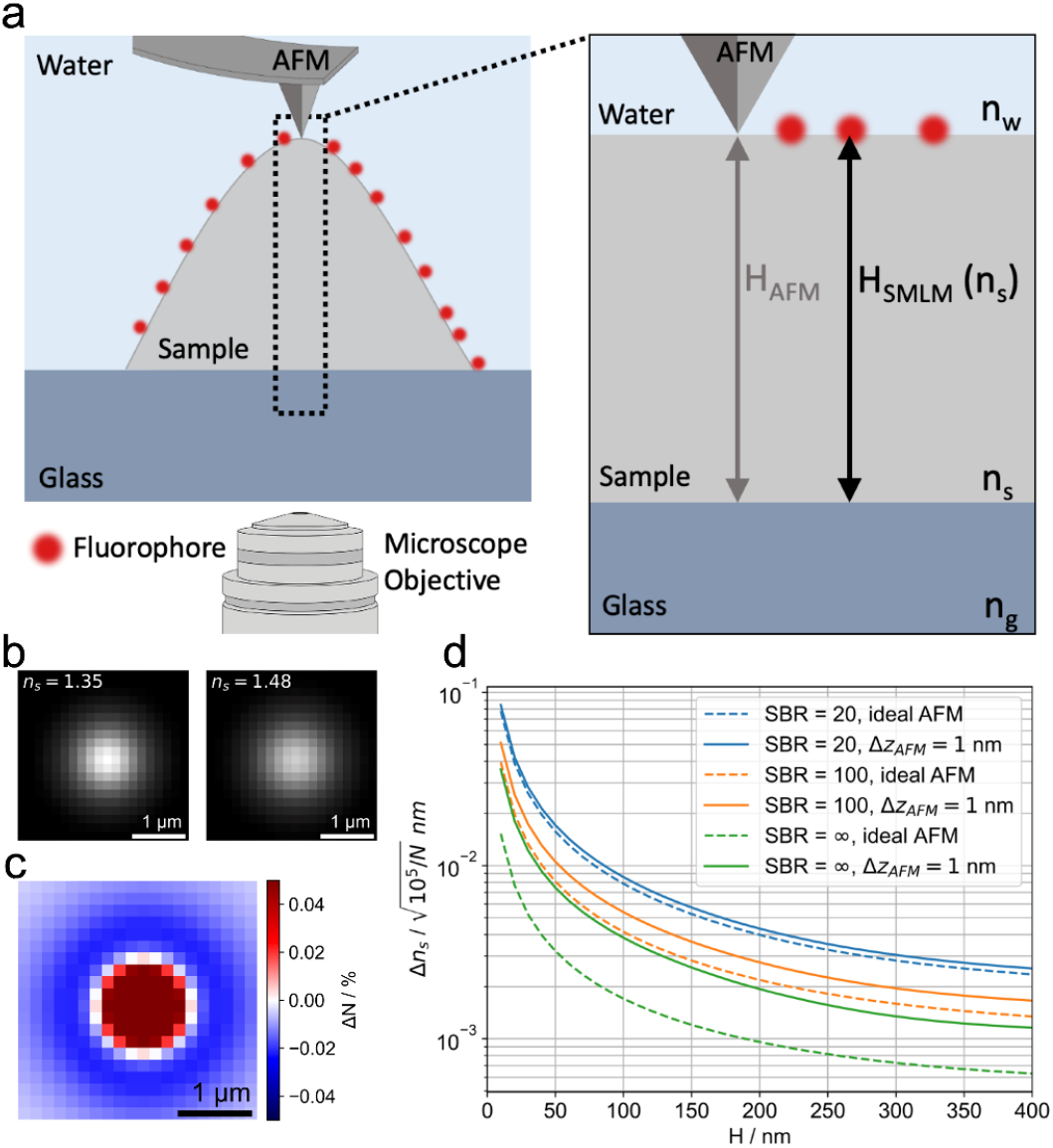
**a** Principle of the refractive index determination method. A biological sample of refractive index n_s_ is surface labelled with dye molecules (red dots). Its apparent height is measured both by SMLM and AFM, yielding H_SMLM_ and H_AFM_, respectively. Due to refraction within the sample, the determined value for H_SMLM_ depends on the assumed refractive index. Comparison with the ground truth value H_AFM_ hence allows calculating the sample refractive index. **b** shows the psf for a freely rotating dye molecule located 100 nm above the glass surface and defocused by 500 nm. The left and right image were calculated for a refractive index n_s_ = 1.35 and n_s_ = 1.48, respectively. The difference image **c** shows the broadening of the psf with increasing n_s_ = 1.35. For the figure we assumed an ambient refractive index of water (n_w_ = 1.33) and a wavelength λ = 670 nm. **d** shows the Cramér-Rao Lower Bound, Δn_s_, for refractive index estimates as a function of collagen thicknesses H. Different colors represent different signal to background ratios as stated in the legend. Dashed lines assume an ideal, error free AFM. Solid lines assume an AFM measurement uncertainty of 1 nm.

First, we were interested in the principal precision for determination of refractive indices, using the new approach. For this we performed a numerical analysis based on the Cramér-Rao Lower Bound (CRLB). **Fig. 1d** visualizes the CRLB for determination of the refractive index, Δ*n*_*s*_, for a sample thickness *H* between 10 nm and 400 nm and different signal to background ratios (SBR). SBR is defined here as the ratio of the signal photons to the mean number of background photons in a single pixel, assuming Poissonian noise. As the precision scales inversely with the square root of the detected signal, CRLB can be adapted to specific measurements by dividing the values by 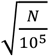, where *N* is the number of collected signal photons in the specific measurement. The plot hence directly reports on Δ*n*_*s*_ for *N* = 10^5^ photons, a value that can be easily obtained with SMLM. As expected, the data indicates that arbitrary precision Δ*n*_*s*_ can be reached, if only the number of detected photons and SBR are sufficiently large. Of note, Δ*n*_*s*_ decreases with increasing sample height *H*, since Δ*n*_*s*_ scales approximately with the relative precision for determining 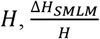 (**Fig. S1**).

For experimental demonstration, we determined the refractive index of isolated single collagen fibrils derived from mouse tail-tendons. Collagen fibrils were immobilized at low density on glass coverslips and fluorescently labelled via AF647-conjugated primary antibody against collagen type I (the most abundant collagen type found in tendon collagen fibrils), giving rather homogeneous surface staining of the fibrils. SMLM imaging was performed at a defocus of appr. 500 nm, which is appropriate for optimal localization precision ^12^. From repeated observations of the same fluorescence molecules, we calculated an axial single molecule localization precision of appr. 5 nm (standard error of the mean of all merged localizations). An overlap of all axial single molecule observations of an exemplary collagen fibril is shown in **Fig. S2a**. For estimation of the fluorophores’ axial positions, we varied the assumed refractive indices of the collagen fibril, which strongly affected the height of the cross-sectional profile. We quantified the fibril height from smoothened data at the crest of the fibril, yielding an apparent height *H*_*SMLM*_(*n*_*s*_) (see Methods for details), which depends on the assumption of the refractive index *n*_*s*_. Next, we recorded the very same collagen fibril regions using AFM (**Fig. S2b**) and determined the ground truth height *H*_*AFM*_. Comparison of *H*_*SMLM*_(*n*_*s*_) with *H*_*AFM*_ allowed for calculating the refractive index of the studied collagen fibril (**Fig. S2c**). For this particular collagen fibril, we obtained *n*_*collagen*_ = 1.437 ± 0.003, which is in good agreement with refractive indices reported for hydrated collagen films ^13^. Here, we determined *H*_*SMLM*_ from a sliding window containing 40 localizations, yielding in total *N* = 123,648 photons contributing to the determined height of the cross-sectional profile. With the obtained *SBR* = 110 and the height of the fibril *H*_*AFM*_ = 138 nm we calculate the theoretical CRLB for the refractive index of Δ*n*_*s*_ = 0.0034, in good agreement with the experimentally obtained value *σ*_*n*_ = 0.0029.

Collagen fibrils swell upon hydration resulting in cross-sectional area increase by up to a factor of four compared to the air-dried state ^14-17^. When submerged in aqueous solution, substantial amount of free (unbound) water fills the intermolecular space within the collagen fibril. Because of the high dielectric constant of water, this reduces the free energy between collagen molecules which manifests as an increase in the intermolecular distance ^18^ and therefore swelling. Consequentially, the degree of swelling also influences the refractive index of collagen, with a convergence towards the refractive index of water for highly swollen collagen ^19,20^. However, current data are only available for average refractive indices obtained from either collagen-rich tissues like cornea ^21^, or from collagen fibers ^20^ and collagen films ^13^, but not at the level of single collagen fibrils.

A number of factors influence the degree of hydration of single collagen fibrils, including cross-linking of neighboring collagen molecules ^22^. Indeed, when comparing the same collagen fibrils recorded in dried and wet conditions, we observed an increase in cross-sectional area varying between 150% and 350%. Accordingly, also the refractive indices obtained for different fibrils varied strongly, ranging from *n*_*collagen*_ = 1.38 up to *n*_*collagen*_ = 1.48 **(Fig. 2a)**. Data from different regions of the same collagen fibril are shown in the same color. When we plotted the calculated refractive index for each collagen fibril versus its area change upon hydration, Δ*A*, we observed a strong negative correlation: in general, the higher the swelling the lower was the refractive index under hydrated conditions. This is in agreement with a model, in which water incorporates into the fibrillar structure upon swelling, resulting in an increase of the intermolecular distance between neighboring collagen molecules ^14,18^; accordingly, *n*_*collagen*_ approaches the refractive index of water for strongly hydrated samples ^19,20^. Still, we observed a substantial spread of the data which exceeded the precision of our method, indicating that the degree of swelling is not the only determinant of collagen refractive index. Even more so, in **Fig. 2a** we indicated in dashed lines the predicted decrease of *n*_*collagen*_ with increased swelling.

**Figure 2:**
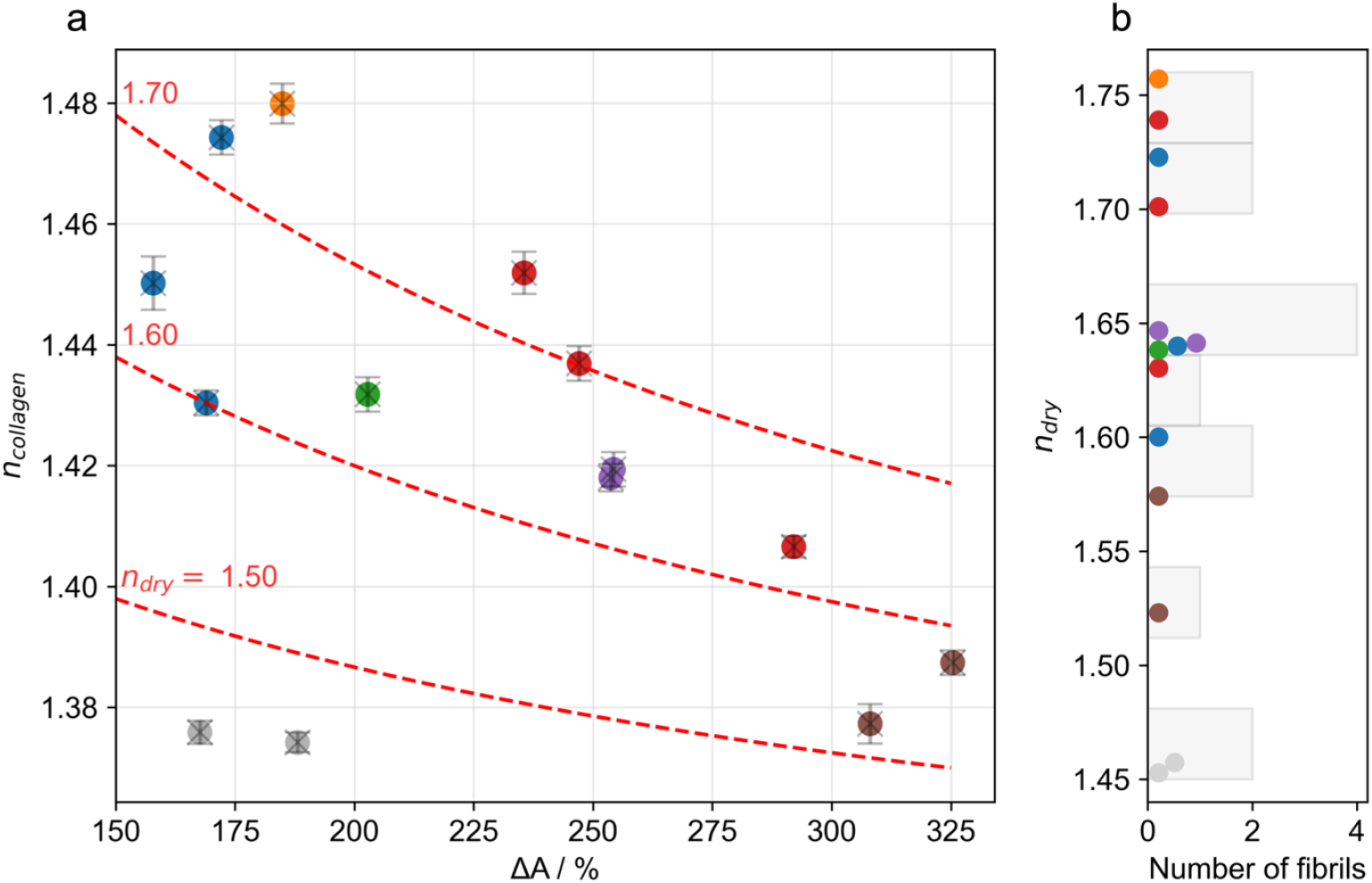
**a** Refractive index of single hydrated collagen fibril, n_collagen_, as a function of the swelling upon rehydration, ΔA. Same colors indicate different regions analyzed on the same fibrils. Red dashed lines show the expected dependence of n_collagen_ on ΔA using Eq. (1), assuming different refractive indices for collagen in the dry state, n_dry_ (red numbers). **b** Histogram of the calculated refractive index for dry collagen, n_dry_, using Eq. (1) for the data shown on the right panel. Data are from 3 independent experiments.

For this, we assumed *n*_*collagen*_ to represent the area-weighted average of the refractive index of dry collagen, *n*_*dry*_, and water, *n*_*water*_ = 1.33:

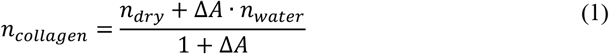

Apparently, this simple model fails to describe the data. We believe the deviation to be a consequence of the violated ergodicity of the collagen sample: evident also from previous work ^17^ different collagen fibrils – albeit from the same source – can be expected to vary in the degree of dehydration that is achieved by drying the sample, potentially due to differences in the amounts of crosslinks between neighboring collagen molecules. As a general trend, smaller amounts of crosslinks will i) lead to larger swelling upon rehydration ^17^, resulting in larger values of Δ*A*, and ii) higher degrees of residual wetting of collagen molecules in the dry state, resulting in smaller values of *n*_*dry*_. In **Fig. 2b**, we showed the estimated spread of *n*_*dry*_, yielding 1.45 ≲ *n*_*dry*_ ≲ 1.75.

Finally, we investigated how the refractive index varied along a single collagen fibril. Indications for refractive index variations were already visible in **Fig. 2a**, apparently, the refractive indices of single collagen fibrils varied more than the according error bars. In **Fig. 3** we show an exemplary collagen fibril (overview, see **Fig.3a**) and the mapped refractive index along the central axis within a sliding window of 100 single molecule signals, corresponding to a spatial resolution of 250 nm in **Fig 3b**. The colors and positions of the dots indicate the determined refractive index and the maximal positions of the corresponding height profiles, respectively.

**Figure 3:**
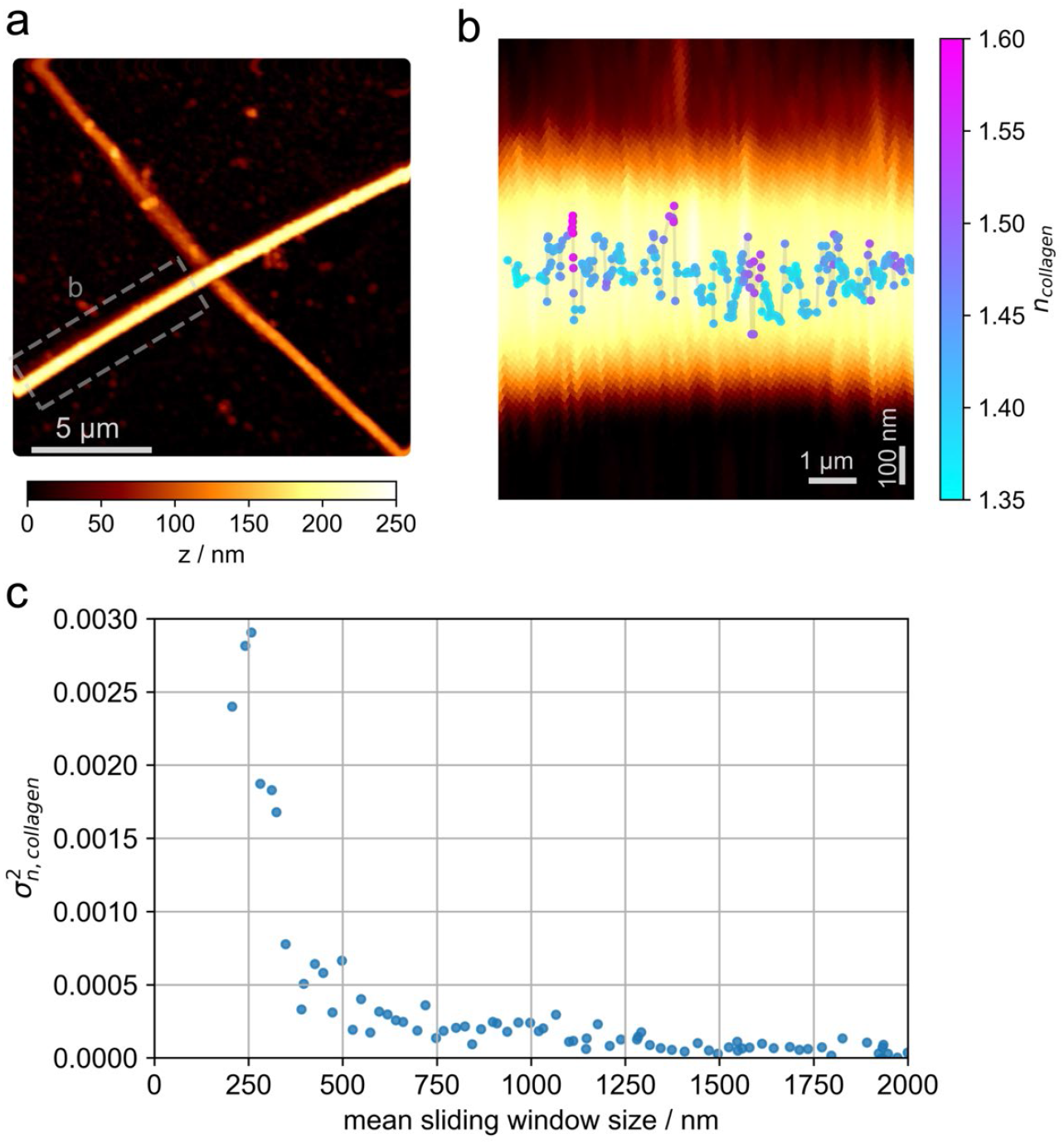
**a** Exemplary AFM image of two collagen fibrils in hydrated conditions. **b** Overlay of the AFM image from the indicated region in **a**, and the refractive index n_collagen_ along the central fibril axis. Each datapoint was calculated from a cross-sectional profile within a sliding window containing 100 localisations, with a 90% overlap between windows; the mean sliding window size was 250 nm. In each window, H_SMLM_ was determined as described in the Methods section. Each dot was plotted at the calculated mean position of all localizations within each window. **c** Variance of the refractive index along the fibril shown in panel **b** for different sizes of the sliding window. Experimentally determined variances, 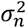, were corrected for experimental errors 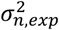 according to 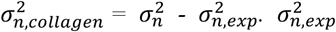 was calculated separately for each sliding window (see Methods section).

Apparently, the refractive index varies substantially in a range between 1.35 and 1.60, indicating differently hydrated regions along the collagen fibril. To study the characteristic length scale of the refractive index fluctuations, we plotted the variance 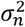 for different sizes of the analysis window. Importantly, both experimental noise, 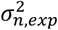, and refractive index fluctuations of the collagen fibril itself, 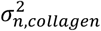, may contribute to the obtained fluctuations. Since the two noise contributions are uncorrelated, however, they can be disentangled and the total variance is given by 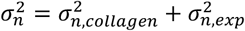. Furthermore, the experimental noise 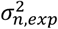 for each sliding window size can be calculated (see Methods, refractive index error estimation), which allows for determining 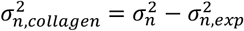 (**Fig. 3c**). For convenience, we show on the x-axis instead of the window size the according mean distance along the collagen fibril. Interestingly, the variance shows a steep transition from large values at distances < 500 nm to rather small values at larger distances. In view of the length of single collagen molecules of 300 nm, our data hence indicates substantial variability in collagen hydration at the length scale of single collagen molecules within collagen fibrils. Similar results are shown for a second fibril in **Fig. S3**.

## Discussion

In this paper we describe a method to extract information on refractive properties of the sample medium from the psf shape. For this, we utilized the possibility to convert an *a priori* ill-conditioned into a well-conditioned problem, employing additional complementary information. Here, we showcase the method by calculating the average refractive index of the sample at high precision, and by mapping the refractive index at lower precision; in our example, the additional information comes from AFM height recordings of the sample. In the following, we describe i) the limitations of our method and ii) its pros and cons compared to other methods for refractive index determination. Finally, in point iii) we discuss our results in the context of collagen properties.

i. *Limitations in the precision for determining n*. In principle, the main limitation of the method is set by the accuracy with which height can be measured. In our example, we demonstrate the method with STORM recordings, yielding an axial localization precision of ∼5 nm. An alternative method would be DNA-PAINT, which can achieve even higher localization precision due to basically unrestricted amounts of localizations that can be merged to infer the position of a fluorescent label ^2^. Next, also the type of labelling needs to be considered. Here, we employed primary antibodies directly conjugated to dye molecules. We referenced the positions of the antibodies at the apex of single collagen fibrils against the positions of antibodies unspecifically bound to the glass coverslip. Reducing the label size would allow for further improving the precision of the method. In addition, one needs to rule out other effects on the psf shape. For example, it has to be ensured that the dye molecules rotate freely, as also azimuth and inclination angle affect the psf shape ^10^. This becomes even more pronounced when analyzing super-critical angle fluorescence ^23^. Here, we used freely rotating dyes conjugated to antibodies in order to avoid orientation biases. Moreover, residual aberrations of the imaging system will affect the psf shape. We corrected for such aberrations in our study using psf recordings of fluorescent beads. Eventually, one may analyze much smaller samples than the chosen collagen fibrils. However, height calculations both from AFM and SMLM show precisions that are almost independent from the actual height of the sample. Therefore, the thinner the sample gets, the higher the relative errors will be, and the more signal photons need to be collected to obtain a given minimum precision for Δ*n*. In **Fig. S4** we show the required number of collected photons for the determination of the refractive index at a precision of Δ*n*_*s*_ = 10^−2^ as a function of sample thickness. We see that in the case of an AFM measurement with an error of 1 nm and a realistic SBR of 100 in SMLM, about 10^5^ photons need to be collected to achieve the defined minimum precision for samples of 50 nm thickness. This corresponds to > 10^2^ localizations at an average signal of 10^3^ photons, which is within realistic limits. The refractive index of samples with a thickness below 40 nm can only be determined to an accuracy of Δ*n*_*s*_ = 10^−2^, if ground truth information on the sample height is known with better precision than 1 nm.
ii. *Alternative methods to determine the refractive index of biological samples at high spatial resolution:* The refractive index affects various optical properties, which can be experimentally assessed ^24^. It is hence not surprising that a variety of methods exist which allow for determination of the refractive indices of biological samples. A common technique is optical coherence tomography (OCT), where differences between the optical and the physical path length are determined and used for calculating *n* ^25^. While this method provides non-invasive insights into the optical properties of samples, its sensitivity and spatial resolution is insufficient to characterize samples with sizes of 100 nm or less. Also ellipsometry is frequently used to determine the refractive index of thin samples ^26^, albeit at classical diffraction-limited resolution. Coupling to a near-field tip can in principle improve the resolution below the diffraction limit, yet the quantitative effect of the tip presence on the determined refractive index is not fully understood yet ^27^. Recently, interferometric detection of scattering (iSCAT) was applied to calculate the refractive index of subdiffraction objects, based on their scattering contrast ^28^. Similar to SMLM also iSCAT contrast depends both on *n* and on the size of the particle *d*, and hence requires an independent measurement of *d*. In ^28^ single particle tracking was used to estimate *d* of the spherical objects based on the measured diffusion constant, using Stokes-Einstein theory.
iii. *Discussion on the obtained values of n*_*collagen*_. Our data on hydrated single collagen fibrils are generally in good agreement with refractive indices, as obtained for various collagen samples ^13,19,21^. For example, Wang and colleagues measured the refractive index of collagen films via OCT and obtained *n* = 1.43 and *n* = 1.53 for hydrated and dehydrated samples, respectively ^13^. A larger spread was observed for fascicles also via OCT, yielding *n* = 1.37 and *n* = 1.57 for hydrated and dehydrated samples ^19^. However, all results reported so far were obtained on rather large collagen samples, where data correspond to an average refractive index over multiple collagen fibrils. This precluded direct correlation between refractive index and swelling behavior upon hydration.

Our high-resolution imaging approach, in contrast, allowed us to correlate the single fibril refractive index to its particular swelling behavior. We found a surprising heterogeneity both in the swelling behavior as well as in the refractive index, potentially due to different amounts of inter-molecular crosslinks within single fibrils ^17,29-31^. From the degree of swelling and the refractive index of water we estimated the expected refractive index of each fibril in the dry state, yielding a spread from *n*_*dry*_ = 1.45 up to *n*_*dry*_ = 1.75. We interpret the rather large variation in refractive index as a consequence of variations in the residual hydration of collagen in the dry state, potentially resulting from variations in the amounts of inter-molecular crosslinks: in this view, a low refractive index would correspond to collagen fibrils with weakly crosslinked collagen molecules, showing higher degree of residual hydration at the air-dried state; a high refractive index, in contrast, would relate to collagen fibrils with well-crosslinked collagen molecules, from which unbound water was efficiently expelled during drying.

We observed this variability in refractive index even at the level of single collagen fibrils, with a strong distance-dependence: *n*_*collagen*_ varies strongly at length scales below 500 nm along the fibril axis and less at larger distances. This behavior may indicate variations in the amount of inter-molecular crosslinks at the level of single collagen molecules. Single collagen molecules are 300 nm in length and twisted into a triple helix. Upon translation, collagen molecules undergo several modifications at the ER, including the enzymatic hydroxylation of lysine. Variability in the degree of lysine hydroxylation between different collagen molecules can hence be expected. Importantly, the presence of hydroxylysine is a prerequisite for the formation of a class of covalent cross-links between adjacent collagen molecules during fibrillogenesis in the extracellular matrix ^32^. Together, the propensity of collagen to form intermolecular cross-links likely varies on the length scale of appr. 300 nm, which would lead to corresponding variabilities in the local hydration and, hence, refractive index.

Finally, it is interesting to compare our results with refractive indices of other proteins, as obtained at different humidities. For green fluorescent proteins, e.g., the refractive index increased from 1.45 to 1.75 from the hydrated to the dry state ^33^, in good agreement with the predictions of our study.

## Methods

### Materials

- Mouse tails (wild type, 6 months old, female; provided by Peter Pietschmann (Medical University of Vienna, Austria)
- PBS (DPBS, D8537-1L, Sigma Aldrich)
- anti-collagen1(I) antibodies, Alexa Fluor 647 conjugated (antibodies, Rabbit, polyclonal, bs-10423R-A647, Bioss)
- BSA (A3803-10G, Sigma Aldrich)
- PFA (stock solution 16% PFA, methanol free, MP Biomedicals)
- glucose, glucose oxidase, catalase, cysteamine (Sigma Aldrich).
- precision tweezers (fine tips, Inox08,5, Dumont)
- coverslip (24×60mm, #1.5, Menzel)
- dental glue (Twinsil extra hard, Picodent)
- Fluorescent beads (FluoSpheres, Cat.Nr. F8780)

### Collagen sample preparation

Collagen fibrils were extracted from frozen mouse tails. The tails were first hydrated with PBS to thaw and then prepared according to our sample preparation procedure, which is sketched in detail in **Fig S5**.

First, the tail was carefully skinned, and the tendons were extracted. Using a scalpel and precision tweezers, the tendons were further separated under a stereo microscope (Zeiss Stemi 508) to expose the collagen fibers. These fibers were then meticulously opened to reveal the collagen fibrils, which were extracted onto a 10 min plasma-cleaned (Plasma Cleaner PDC-002 (230V) from Harrick) glass coverslip. After preparing the samples on the coverslips, the samples were rigorously washed with deionized water for at least 30 seconds to remove excess collagen material and salt residues. The prepared samples were then dried using nitrogen gas and stored in an air-tight chamber until the experiment.

### Antibody labelling

Prior to antibody labeling of the collagen fibrils, a custom-made fluid cell was mounted onto the coverslips using two-component dental glue. The fluid cell was specifically designed with wells large enough to accommodate the AFM glass block holding the cantilever, while featuring sufficiently high walls to prevent any liquid spillage during measurements, ensuring a controlled and stable environment for imaging. Once the fluid cell was in place, a small scratch was made on the underside of the coverslip, serving as a positional reference marker, enabling accurate alignment when transferring the samples between the different setups.

Next, the collagen fibrils were labelled for one hour at room temperature, using primary antibodies conjugated with Alexa Fluor 647. For labelling, we diluted anti-collagen1(I) antibodies 1:1000 in PBS containing 1% BSA. After labelling, the staining solution was removed and the samples were carefully rinsed with deionized water to clear the samples from excess dye. The samples were then fixed at room temperature for 15 minutes, using 4% paraformaldehyde (PFA) diluted in deionized water. Following fixation, the PFA solution was removed and the samples were thoroughly rinsed with deionized water to eliminate any residual PFA. Finally, PBS was carefully applied to the samples to maintain fibril hydration prior to imaging.

### Single molecule localization microscopy

A home-built experimental setup was used as described previously ^8^. It is based on an Olympus IX73 (Japan) microscope body equipped with a high NA objective (Carl Zeiss, alpha-plan apochromat, 1.46 NA, 100x, Germany) and an EMCCD camera (Andor iXon Ultra). Furthermore, it features a red excitation laser (640 nm laser light, 100 mW nominal laser power, OBWAS Laser box, Coherent, USA). The setup was operated in spinning TIRF excitation (iLas^2^), yielding an excitation intensity of 1 kW/cm^2^. Lasers were filtered by a quad dichroic mirror (Di01-R405/488/532/635, Semrock, USA) and an emission filter (ZET405/488/532/642m, Chroma, USA) placed in the upper deck of the microscope body. Illumination and image acquisition was operated by VisiView (Visitron Systems, Germany). To maintain constant focus during imaging, a focus-hold system was built based on an IR laser, which was totally reflected at the Surface of the coverslip. The reflected beam was then captured by an IR camera. Any vertical (z-axis) movement of the sample causes a lateral shift in the beam on the IR camera chip ^34^. A z-piezo underneath the objective was used both for focus-holding and for deliberate defocusing the sample, as required for determining the z-positions of the single molecule signals (see subsection “Data analysis for single molecule localization microscopy” below).

Fluorescent beads were added to the sample to serve as fiducial markers. The beads were sonicated in an ultrasonic bath for 2 minutes, diluted 1:100 000 in PBS and finally applied to the sample for 5 min. Afterwards the sample was rinsed with PBS. For dSTORM imaging ^35^, PBS was exchanged with blinking buffer consisting of: 10% glucose, 500 μg/ml glucose oxidase, 40 μg/ml catalase and 50 mM cysteamine in PBS (pH 7.5). The dSTORM buffer was always prepared fresh immediately prior to imaging or replaced between measurements, as its pH tends to shift after one hour, potentially negatively affecting imaging quality ^36^.

For dSTORM imaging, the first step was to select a suitable region of interest (ROI). Each ROI contained at least two collagen fibrils featuring an intersection, but were free of excessive overlapping fibrils. While overlapping fibrils can theoretically be imaged using STORM, they usually have heights that exceed the TIRF excitation range and are more susceptible to movement, which renders them unsuitable for the imaging process. Any overlapping fibrils were therefore excluded from further analysis. After identifying a ROI, the x and y coordinates of the microscope stage were recorded, together with the reference marker, to enable precise relocation of the same ROI during subsequent AFM measurements. Next, the fluorophores within the selected region of interest (ROI) were excited in epifluorescence mode using the red laser until the majority entered a dark state, demonstrating blinking behavior. For dSTORM imaging, the illumination time was set to 20 ms and the camera delay was adjusted to 21 ms, the excitation was switched to TIRF mode, and the camera gain was adjusted to 300.

The intensity of the red laser was maintained at 1 kW/cm^2^, while the intensity of the UV laser (wavelength of 405 nm) was set to 5% of the nominal power of 100 mW. The UV laser was used to excite the fluorescent beads used as fiducial markers every 100^th^ recorded frame. Additionally, the UV laser aided in reactivating the fluorophores on the collagen, allowing for extended STORM recording durations ^1^. The sample was then defocused, typically 500 nm above the coverslip unless specified otherwise. At least 80,000 images were recorded for each ROI; if the imaging time of a sample exceeded 50 minutes, the STORM buffer was exchanged for a fresh solution. Furthermore, we obtained the aberrations from the setup by measuring the psf from a z-stack of fluorescent beads with the same setup configuration that was used for the dSTORM measurements.

### Atomic force microscopy

After STORM imaging, the samples were washed, dried and transferred to the AFM setup. The ROIs were relocated, using the reference marker (glass scratch) and the recorded microscope stage positions. The samples were then imaged with the AFM, first under dry condition in AC-mode and afterwards the samples were (re-)hydrated with PBS and imaged in Qi-mode.

An AFM from JPK (Nanowizard 3, Bruker, Germany) was used with the following tips: i) for measuring in AC-mode: PPP-NCHR-50 (from Nanosensors), width: 30 μm, length: 125 μm, thickness: 4 μm, frequency: 330 kHz, spring constant: 42 N/m, tip radius 7 nm; ii) for recordings in liquid using the Qi-mode: MSCT-F (Bruker AFM Probes), width: 18 μm, length: 85 μm, thickness: 0.6 μm, frequency: 125 kHz, spring constant: 0.6 N/m, nominal tip radius: 10 nm and typically a setpoint of 1.5 nN was chosen.

AFM images with a size of 20 μm x 20 μm were recorded to ensure sufficient information for comparison with the dSTORM data, which has a usable image size of approximately 30 μm x 30 μm. The camera connected to the AFM setup was a DMK 31BF03 Monochrome Camera and the used objective for the AFM measurements was an air objective, Zeiss Plan-Neofluar 20x/0.5, 440340.

### Data analysis for single molecule localization microscopy

The aberrations of the setup were calculated from the recorded bead data, using the Matlab application Aberration Measurement from A. Jesacher et.al.. For the localization of the fluorophores, the python application mlefitgpu (https://github.com/jgmaloberti/mlefitgpu) was used. The software allows for pre-localization of the fluorophores, as well as modelling and fitting of psfs in order to determine the position of the fluorophores in 3D.

Fluorophores were pre-localized using their intensity values (absolute and relative pixel intensity). The PSF was modelled using the imaging setup details and the calculated aberrations. The PSF model allows for several input parameters, including the defocus, middle layer thickness, refractive index of the used media surrounding the fluorophores and the refractive index of the middle layer (collagen).

To find the exact defocus value of a single region of interest (ROI), a reference plane was first defined, using only the fluorophores attached to the coverslip, adjacent to the collagen fibrils. Python codes for analysis are available under the following link: https://github.com/simonjaritz/SMLM-Analysis.git. The refractive index of the surrounding media was chosen to be nw = 1.33 (water). The defocus is determined by setting the reference plane to z = 0 nm. Afterwards, the height of the same fibril was determined from the AFM data, where the hydrated samples were imaged in Qi-mode.

Using the determined defocus value and the reference height from the AFM we re-analyzed the fibril using now an additional middle layer in our PSF-model. For the analysis of one fibril, we kept the height of the additional layer constant (fibril height from AFM), but varied its refractive index between 1.38 and 1.48. The individual fibrils were then further analyzed to obtain the cross-sections and calculate their heights, which were then matched with the AFM data.

### Analysis of collagen cross-sections

The single molecule localizations were corrected for lateral drift using the recorded fiducial markers and tilt corrected, using the reference plane at their determined defocus value. The localizations were further filtered, keeping only localizations with high fitting quality or low Log-Likelihood ratio (LLR < 600, with each localization featuring 15×15 pixels). The AFM images were first tilt corrected for each line. To find the contact point of the AFM tip on the collagen fibrils, force distance curves of the Qi-measurements were fitted by a Hertz model by our in-house developed software (https://github.com/Rufman91/ForceMapAnalysis).

Both dSTORM and AFM measurements were further analyzed in order to show all localizations of a fibril in a single cross section. This analysis accounts for the curvature of fibrils. First, the fibrils were fitted with a fourth-order spline (with z = 0) through their central axis and then the shortest distance between the spline and each localization was calculated. All localizations were transformed to new coordinates denoted s, X and Z, for the longitudinal, lateral and vertical axis with respect to the fitted spline. The resulting cross-sections feature all localizations along the fibril and were used for the fibrils’ height determination.

To calculate *H*_*SMLM*_ and *H*_*AFM*_, the localizations were cluster-filtered using the DBSCAN algorithm (available on https://scikit-learn.org/stable/modules/generated/sklearn.cluster.DBSCAN.html), to remove outliers. Next, the median height was calculated within a sliding window. The number of localizations per window was kept constant for each fibril, and the highest binned datapoint was taken as the height of the fibril. All steps of a single fibril analysis measured with SMLM are provided in **Fig. S6**. For determination of *H*_*SMLM*_, we further considered the precise location of fluorophores next to collagen fibrils, which defined the reference plane. The median height of debris at the coverslip surface was determined by AFM and added to *H*_*SMLM*_.

This process was repeated for each modelled refractive index and the heights were then plotted against the refractive index together with the AFM height of a single fibril (see **Fig. S2c)**. Next, the intersection of a linear fit through the dSTORM datapoints and the AFM data was calculated, representing the refractive index of the fibril. Furthermore, the cross-section areas were calculated using the same sliding window binning method for AFM dry, *A*_*dry*_, and AFM wet, *A*_*wet*_, using numerical integration. Afterwards, the refractive index was plotted against the swelling Δ*A*, defined as the relative cross section area increase 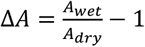.

### Refractive index error estimation

We used resampling to estimate the error of the height calculations *H*_*SMLM*_ and *H*_*AFM*_. Briefly, half the localizations of a single cross-section were randomly selected and the height was calculated for the selected datapoints. This process was repeated 1,000 times and the final standard deviation of the resulting Gaussian distribution was divided by 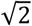, yielding an error estimate for the collagen height.

For the error estimation of the refractive index, we employed random sampling from the obtained Gaussian distributions for height estimates. This step was repeated 10,000 times to obtain the variance for the refractive index estimates 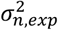.

### Single molecule localization precision

To calculate the localization precision, we tracked blinking signals of one and the same fluorophores over several frames, calculated the standard deviation of the determined positions and used the standard error of the mean as localization precision of the merged signals.

### Determination of the refractive index measurement precision

The statistical error of the refractive index measurement Δ*n* depends on the errors of the SMLM and AFM measurements. Because these errors are unrelated, the respective variances add up:

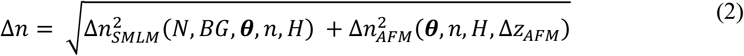

The error contribution of the SMLM measurement, Δ*n*_*SMLM*_, can be derived using a Cramér-Rao Lower Bound (CRLB) analysis. While in SMLM, calculations of the CRLB are usually utilized to determine the localization precision, we follow an analogous analysis here to quantify the best precision to which the refractive index of the sample can be determined by any unbiased estimator, in case we know its true height *H*. For this we defined a function *I*_*x*,*y*_(*N, BG, θ, n, H*) that calculates images of single fluorescent molecules, i.e. numbers of photons detected in camera pixels with indices *x, y*. The function depends on the fluorescence signal *N* and the background level *BG*, several microscope related parameters such as the peak emission wavelength, numerical aperture, camera properties etc., which are contained in a parameter vector *θ*, and finally the refractive index *n* and height *H* of the sample.

The minimal error of the refractive index estimation caused by SMLM is then derived as the square root of the inverse Fisher information

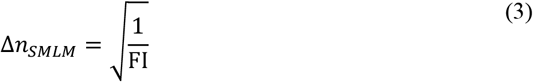

with the Fisher information calculated as

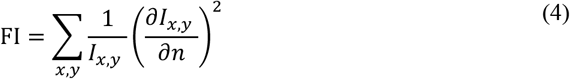

Here we don’t show the explicit dependencies of FI and *I*_*x*,*y*_ on the many arguments for clarity.

To determine the influence of the AFM measurement error Δ*z*_*AFM*_ on the error of the refractive index measurement, Δ*n*_*AFM*_, we calculated (noise-free) molecule images *I*_*x*,*y*_ assuming sample thicknesses *H* ± Δ*z*_*AFM*_, and identified those refractive index values that provide best-matching (noise-free) images under the assumption that the sample thickness is *H*. More precisely we performed the following optimization:

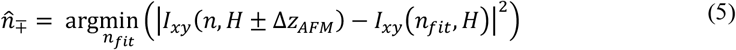

From this, we approximated a symmetric error Δ*n*_*AFM*_ as

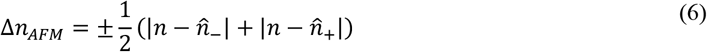

which was feasible since 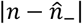 and 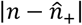 showed only negligible differences.

## Acknowledgements

The study was supported by the Vienna Science and Technology Fund (WWTF) [10.47379/LS19035] and by the Austrian Science Fund (FWF) [10.55776/P36022]. For open access purposes, the author has applied a CC BY public copyright license to any author-accepted manuscript version arising from this submission. The authors would like to acknowledge Peter Pietschmann (Medical University of Vienna, Austria) for his kind provision of mouse tails.

## Author contributions

SJ: performed SMLM/AFM experiments and analysis; LV: performed SMLM experiments and analysis; AG: performed SMLM experiments and analysis; MR: performed AFM experiments and analysis; PJT: conceived collagen experiments and contributed to the discussion of the results; OGA: conceived collagen experiments and contributed to the discussion of the results; JGM: contributed to the SMLM analysis; SM: contributed to the SMLM analysis; AJ: contributed to the SMLM analysis; GJS: conceived the study, contributed to data analysis, wrote the paper

## Supplementary Figures

**Figure S1:**
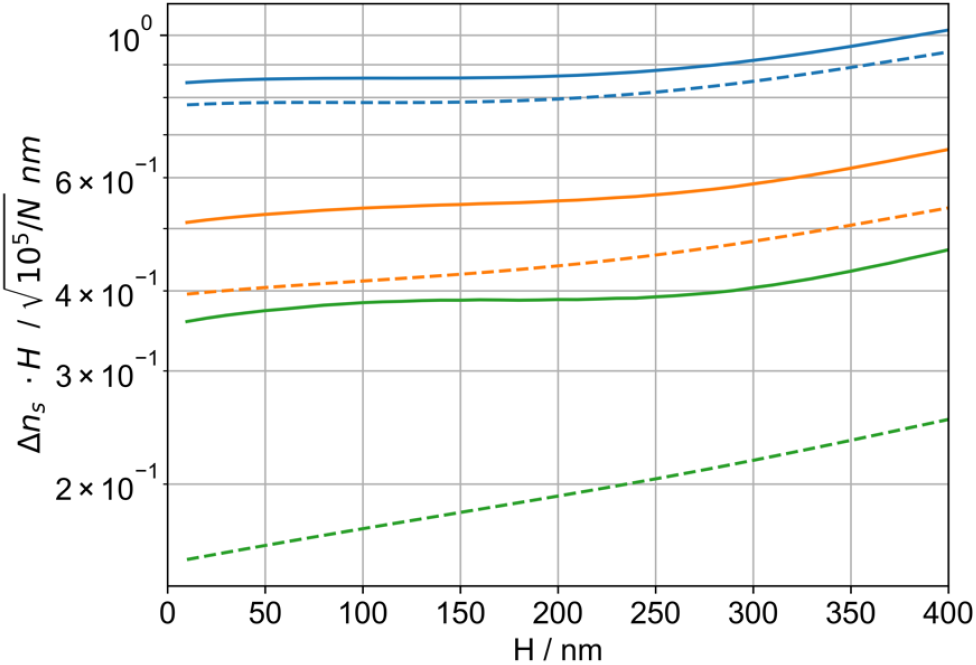
The product of Δ*n* · *H* depends only weakly on the collagen thickness. The plot was calculated for the same parameter settings as **Fig.1d**, i.e. a refractive index of *n* = 1.42 and a signal of 10^5^ photons. Color code is identical to **Fig.1d**.

**Figure S2:**
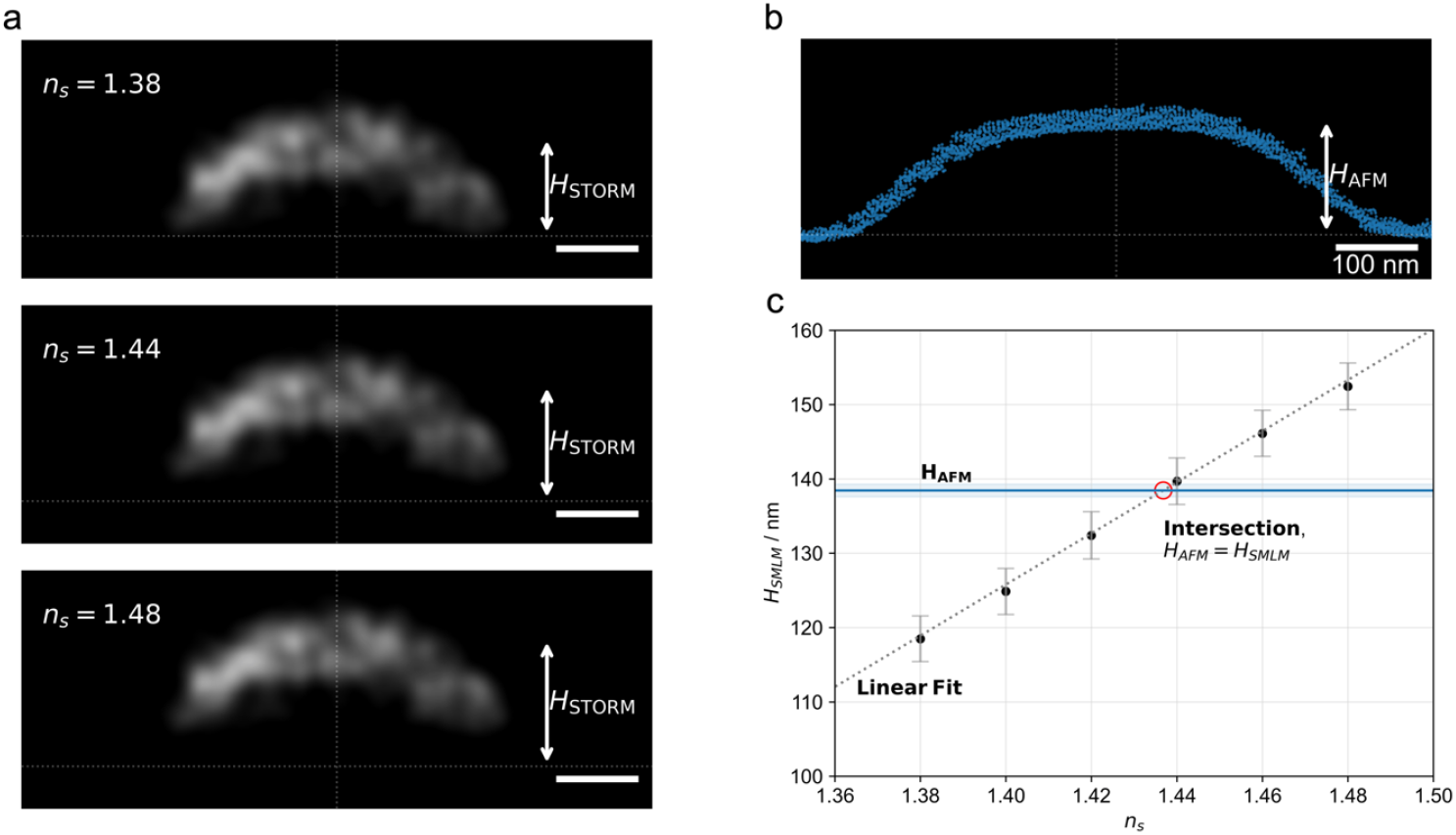
Example of a collagen fibril cross section, imaged with **a** SMLM and **b** AFM in hydrated state. SMLM data were analysed with different assumptions about the refractive index, shown here exemplarily for *n*_*s*_ = 1.38, 1.44, 1.48. The AFM profile yielded *H*_*AFM*_ = 138 nm. Scale bar = 100 nm. **c** Comparison between the ground truth height measured via AFM (*H*_*AFM*_,) and the apparent height determined via SMLM (*H*_*SMLM*_) as a function of the assumed refractive index (*n*). From the intersection, the correct refractive index of the collagen fibril can be determined. Shown are data obtained from a single collagen fibril. For this fibril we calculated a refractive index *n*_*collagen*_ = 1.437 ± 0.003.

**Figure S3:**
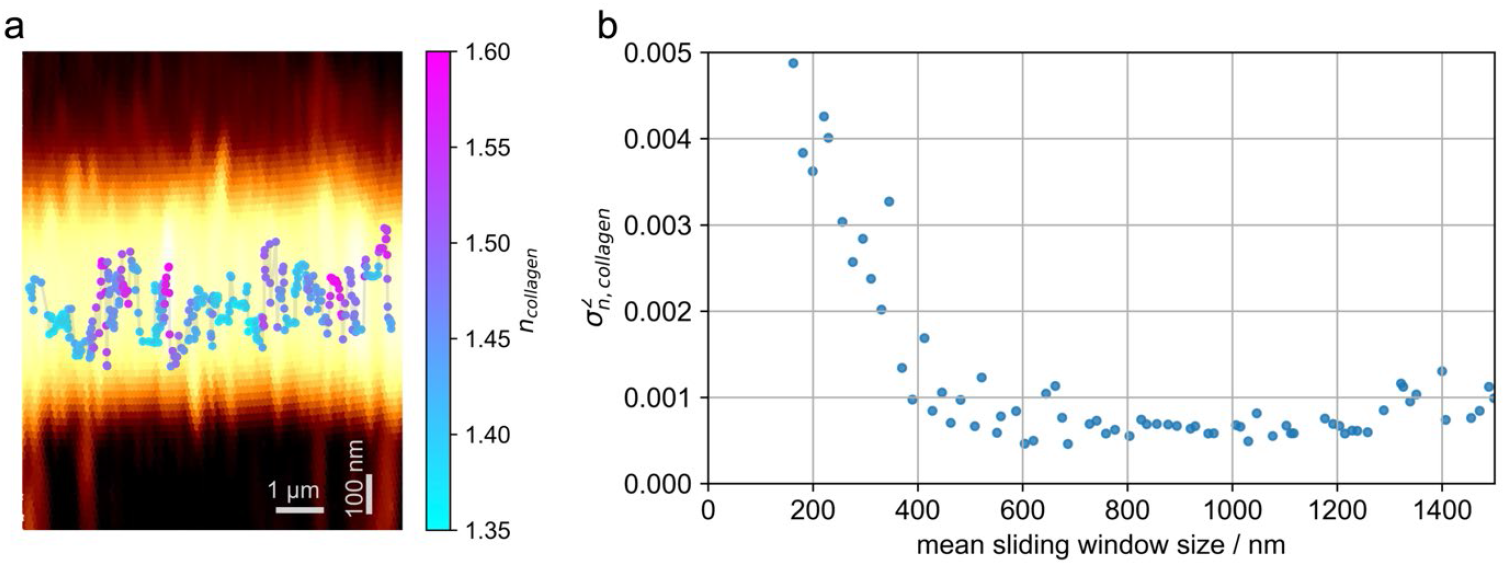
**a** Overlay of the AFM image from an example fibril and the refractive index *n*_*collagen*_ along the central fibril axis. Each datapoint was calculated from a cross-sectional profile within a sliding window containing 100 localisations, with a 90% overlap between windows; the mean sliding window size was 200 nm. In each window, *H*_*SMLM*_ was determined as described in the Methods section. Each dot was plotted at the calculated mean position of all localizations within each window. **b** Variance of the refractive index along the fibril shown in panel **a** for different sizes of the sliding window. Experimentally determined variances, 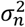, were corrected for the expected errors in determining the refractive index, 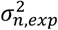, according to 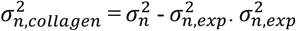 was calculated separately for each sliding window (see Methods section).

**Figure S4:**
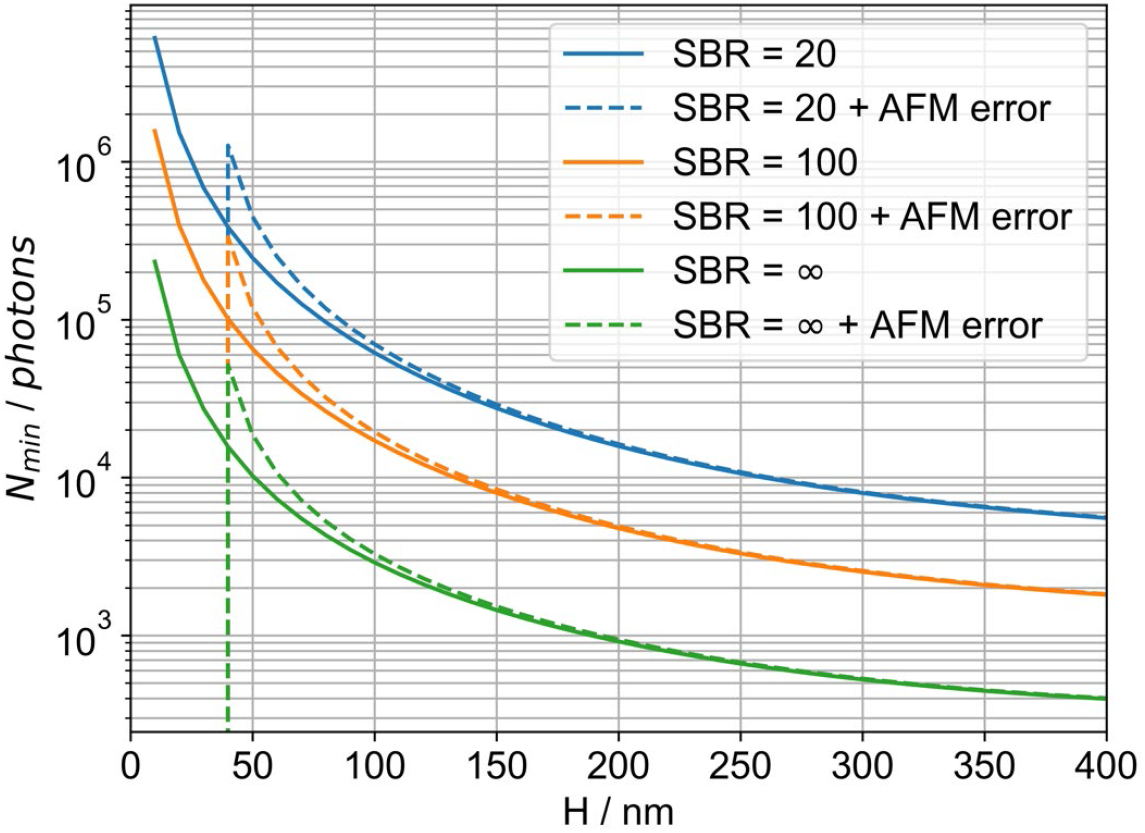
Required numbers of signal photons *N*_*min*_ as a function of sample thickness *H*, to ensure a refractive index precision of Δ*n*_*s*_ = 10^−2^ assuming an ideal AFM (solid lines) and an AFM error of 1 nm (dashed lines).

**Figure S5:**
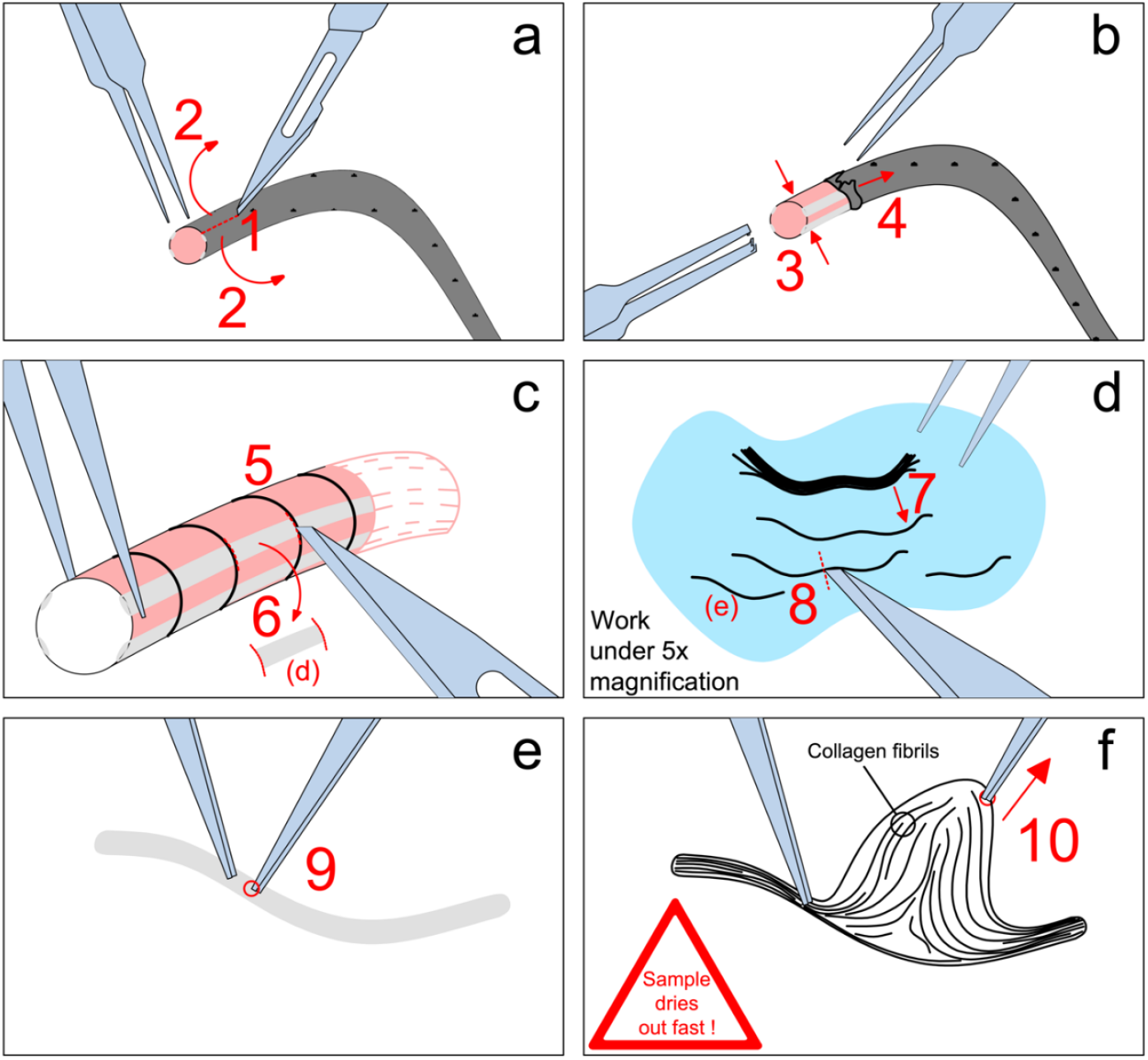
Collagen fibril sample preparation. **a** Cut lengthwise (approximately 1 cm) with a scalpel (1) and pull the skin towards the end of the tail using tweezers (2). **b** Hold the proximal end of the tail tightly with the toothed, thumbed forceps (3) and pull the skin in the distal direction using either tweezers or an additional pair of toothed, thumbed forceps (4). **c** Make perpendicular cuts in the tendons, approximately 1 cm apart (5). Be careful not to cut the muscle tissue underneath and to the sides of the tendons. Next, remove the cut tendon pieces with tweezers and a scalpel (6). Those tendon pieces can then be stored in the freezer till they are used for fibril sample preparation (step **d**) and onwards. Alternatively, a clamp with three teeth can be used to fixate the tail in place and the tendon can be cut out. The cutting is performed utilizing a scalpel and a pair of tweezers similarly as described before. **d** Place a piece of tendon into a round petri dish (Ø 3.5 cm) and hydrate with PBS or sterile water. Place the sample under the stereo microscope using 5X magnification. Use a pair of tweezers to separate the fibres (7) and the scalpel to cut them (8), to get a higher sample yield. Each separated fibre can then be used for the next step (9). **e** Take a piece of fibre and place it onto a 10-minute plasma cleaned coverslip. Next, pinch it with the tip of precision tweezers (9). **f** Pull gently on the rest of the fibril with a second pair of precision tweezers to reveal the collagen fibrils (10). This step has to be performed fast (within a few seconds) as the fibres are prone to drying out.

**Figure S6:**
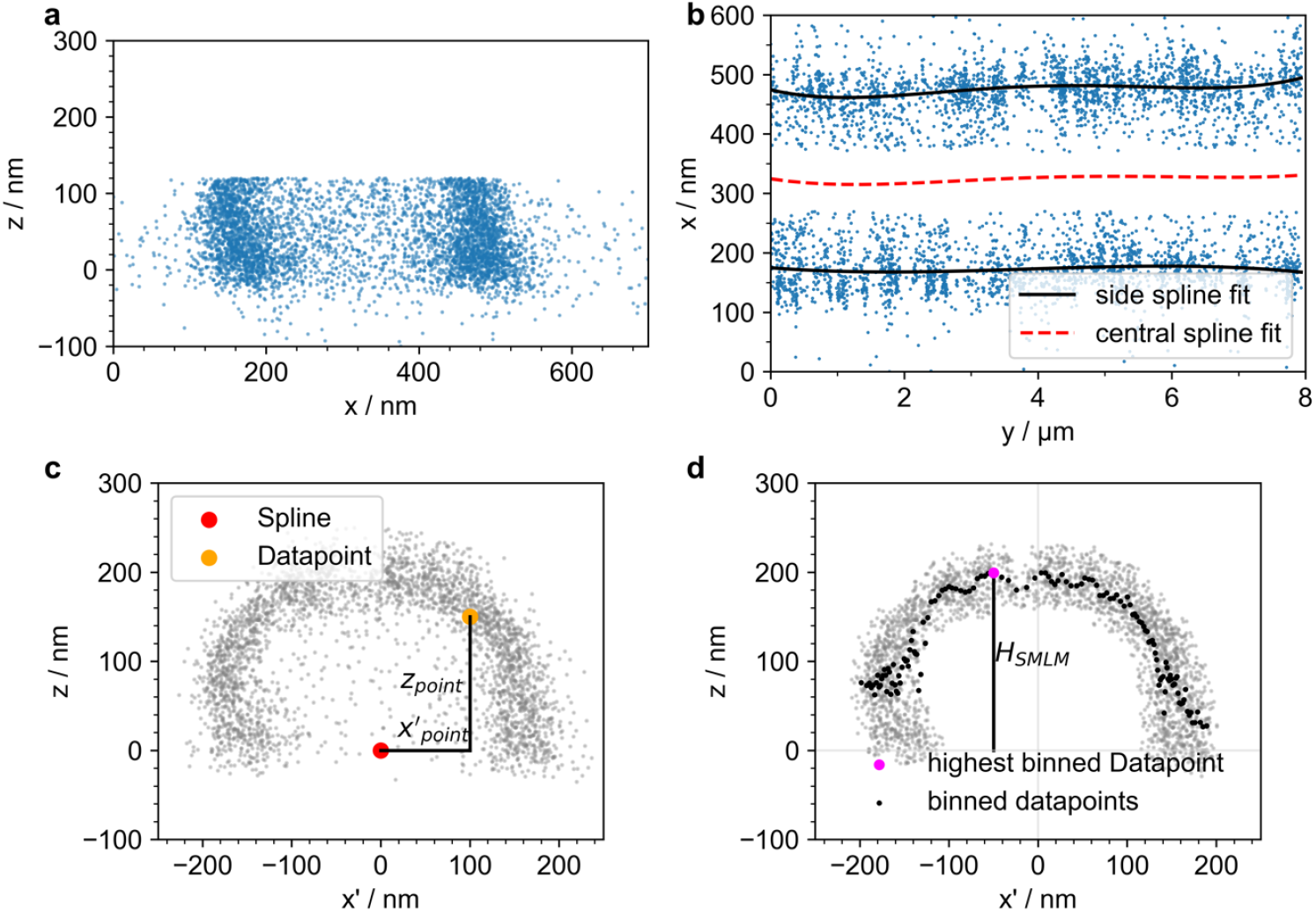
Procedure for height determination of collagen fibrils imaged with SMLM. We first rotated the fibril such that its long axis aligned roughly with the y-axis of the coordinate system. The z-axis was chosen in the vertical direction. **a** Cross-sectional profile of a single fibril projected onto the (*x, z*) plane. Data were cropped at half maximum to highlight the fibril rims. **b** Top view of the same fibril. We selected the two rims of the fibril and fitted the data with 4^th^ order 2-dimensional splines. Next, a central spline was fitted through the two rim splines, which was taken as the collagen as the collagen axis for subsequent analysis. This procedure ensured that localization clusters due to overcounting of single molecule signals had little impact on the fitted central spline. **c** Transformed cross-sectional profile of the fibril. Datapoint coordinates were transformed, with x’ denoting the horizontal distance from the central spline. **d** To determine the fibril height from the transformed cross-section, we removed outliers using DBSCAN and binned the datapoints along the x’-axis using a sliding window. In this example, each window contained 40 localizations and featured an overlap of 50%. Binned data points are shown in black. For determination of the height, *H*_*SMLM*_, we used the highest binned datapoint (magenta).

